# Hydrodynamic stress and phenotypic plasticity of the zebrafish regenerating fin

**DOI:** 10.1101/2021.01.25.428094

**Authors:** Paule Dagenais, Simon Blanchoud, David Pury, Catherine Pfefferli, Tinri Aegerter-Wilmsen, Christof M. Aegerter, Anna Jaźwińska

**Affiliations:** Physik-Institut, University of Zurich, Winterthurerstrasse 190, 8057 Zurich, Switzerland; Department of Biology, University of Fribourg, Chemin du Musée 10, 1700 Fribourg, Switzerland; Department of Molecular Life Sciences, University of Zurich, Winterthurerstrasse 190, 8057 Zurich, Switzerland

**Keywords:** ray bifurcation, skeletal morphogenesis, growth control, biomechanics, viscous shear stress, biomimetic hydrofoils

## Abstract

Understanding how extrinsic factors modulate genetically encoded information to produce a specific phenotype is of prime scientific interest. In particular, the feedback mechanism between abiotic forces and locomotory organs during morphogenesis to achieve efficient movement is a highly relevant example of such modulation. The study of this developmental process can provide unique insights on the transduction of cues at the interface between physics and biology. Here, we take advantage of the natural ability of adult zebrafish to regenerate their amputated fins to assess its morphogenic plasticity upon external modulations. Using a variety of surgical and chemical treatments, we are able to induce phenotypic responses to the structure of the fin. In particular, fin cleft depth and the bifurcation of the bony rays are modulated by the surface area of the stump. To dissect the role of mechanotransduction in this process, we investigate the patterns of hydrodynamic forces acting on the surface of a zebrafish fin during regeneration by using particle tracking velocimetry on a range of biomimetic hydrofoils. This experimental approach enables us to quantitatively compare hydrodynamic stress distributions over flapping fins of varying sizes and shapes. As a result, viscous shear stress acting on the tip of the fin and the resulting internal tension are proposed as suitable signals for guiding the regulation of ray growth dynamics and branching pattern. Our findings suggest that mechanical forces are involved in the fine-tuning of the locomotory organ during fin morphogenesis.

## Introduction

How genetic information encodes a body shape is a century old scientific question (Johannsen, 1911). Since then, a tremendous amount of knowledge has been gathered on the role of genes and epigenetics in guiding the morphogenetic programs (Ecker et al., 2018; Orgogozo et al., 2015). In addition, the impact of physics in the modulation of cell shape and tissue morphogenesis has progressively been recognized (Aegerter-Wilmsen et al., 2007; Buchmann et al., 2014; Heisenberg and Bellaiche, 2013; Vogel, 2013). However, continuous analysis of mechanotransduction during the development of complex vertebrate organs remains very challenging, thus limiting the progress of this research area. A prime example of physical feedback on organ architecture is the adaptation of bone to altered load (Wolff, 1986). The locomotory organs, either on land or in water, are particularly exposed to external modulations, because they interface with forces and movements. Yet, it remains unknown to which extent morphogenesis can be influenced by the mechanical environment.

Among fish locomotory appendages, the caudal fin is the major component of mobility, which generates propulsion power and sophisticated maneuverability (Flammang and Lauder, 2009). One of the most advantageous feature of this appendage is its robust regenerative capacity throughout the entire fish life. After amputation, the caudal fin can efficiently regrow its missing tissues within a few weeks (Pfefferli and Jaźwińska, 2015; Sehring and Weidinger, 2020). The caudal fin is also experimentally convenient, as it is easily accessible for surgery, imaging and morphometric analysis during the regrowth process. Given its major contribution to swimming, the caudal fin is particularly susceptible to hydrodynamic forces at the time of regeneration. Overall, this appendage thus provides an ideal experimental model to study the interface between physics and biology.

The zebrafish *Danio rerio* is the most common fish species in biomedical research, including orthopedics (Busse et al., 2020). Important advantages of this organism are a powerful set of genetic tools and an efficient maintenance in a laboratory facility. In this species, the caudal fin has a fan-like shape with a superficially symmetrical dorsal and ventral lobe, and a central cleft (Fig. 1A, A’). The surface is supported by 16 to 19 principal rays of segmented bones, spanned by soft interray tissue (Marí-Beffa and Murciano, 2010; Murciano et al., 2007; Suniaga et al., 2018). Bones can bifurcate one to three times, so that the space between adjacent bones is nearly constant throughout the fin surface (Fig. 1A, A’, B, B’). This regularity of rays is thought to provide an optimal rigidity of the appendage during swimming. Together, the bone matrix, surface geometry and bifurcation pattern contribute to the biomechanical properties of this main locomotory appendage (Shoele and Zhu, 2009).

**Figure 1.**
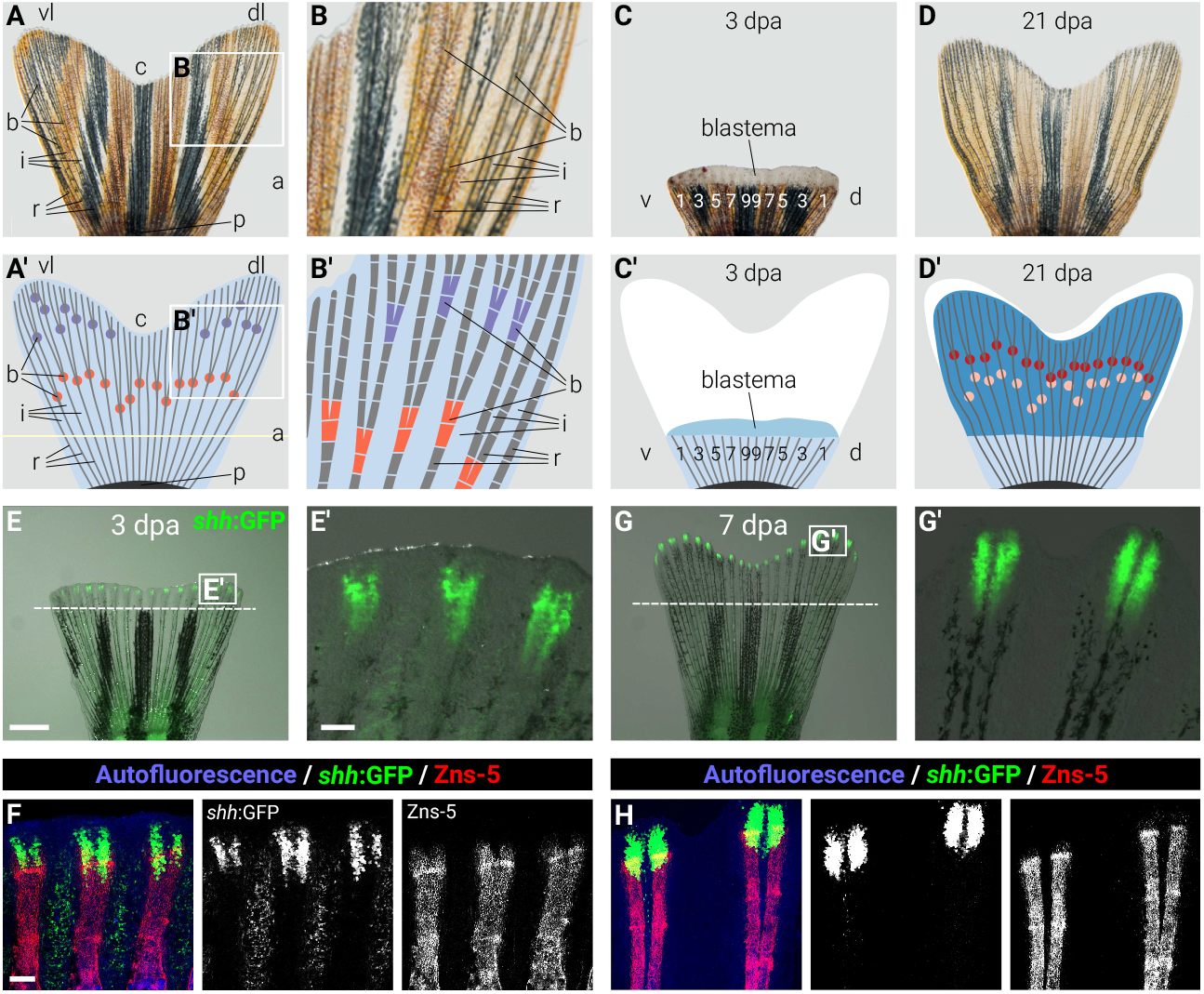
Zebrafish caudal fin morphology and regeneration. **(A-A’)** An intact zebrafish caudal fin. (**A**) Live-imaging of an intact caudal fin of adult zebrafish. The pigmentation stripes have a black and golden appearance and make the fin less transparent. Scale bar is 1 mm. (**A’**) Schematic representation of the same fin displaying the arrangement of bony elements (dark gray lines) and the fin surface (light blue). In this and subsequent figures, the fins are oriented with the distal side to the top, the proximal one to the bottom, their ventral lobe (vl) to the left and their dorsal one (dl) to the right. Relevant structures of the bilobed fin are indicated by lower-case letters and include cleft (c), ray (r), interray tissue (i), ray bifurcation (b) and peduncle (p) at the base of the fin. The amputation plane (a) of the subsequent surgery is depicted with a yellow line. Primary ray bifurcations are highlighted in orange and secondary ones in purple. (**B**-**B’**) Magnification of the areas delimited by the white rectangle in **A** and **A’**, respectively. The bony rays (gray rods) are regularly segmented (white interruptions). The ray bifurcation sites are the regions where the rays split into two symmetrical structures. (**C**-**C’**) Live imaging and schematic drawing of the stump at 3 days post amputation (dpa). The unpigmented outgrowth emerges distally to the amputation plane that contains a growth zone, called the blastema. **(C’)** The shape of the missing part of the fin is depicted in white, and the blastemal outgrowth in darker shade of blue. Indexes of ventral (v) and dorsal (d) rays are overlaid on the corresponding elements of the stump. The fin comprises approx. 8 ventral and 8 dorsal rays, labeled as v1-v8, and d1-d8, respectively. (**D**-**D’**) Regenerated caudal fin at 21 dpa. The new tissue is depicted in dark blue. New primary bifurcations are highlighted in red. A distal shift is observed when compared to the original position of the corresponding bifurcations prior to injury (light orange). (**E-F**) Early detection of bifurcation events by visualizing *shh* expression at 3 dpa. (**E**) Overlaid fluorescence and bright-light imaging of *shh:GFP* transgenic fish. *shh:GFP* (green) is expressed within the blastema at the tip of each ray. Scale bar is 1 mm. (**E’**) Higher magnifications of the area delimited by the white rectangle in **E**. Splitting of the *shh:GFP*-expression region precedes the bifurcation event. Scale bar is 100 μm. (**F**) Confocal images of fixed fins immunostained with Zns5 antibodies (red) that detect osteoblasts, GFP antibody (green) and tissue autofluorescence (blue). Distal-most osteoblasts follow the direction of the *shh:GFP*-expressing cells. Scale bar is 100 μm. (**G-H**) Bifurcated rays at 7 dpa, depicted similarly as in **E-F**.

Surgical amputation of the caudal fin (Fig. 1C, C’) induces a regenerative process that will completely restore this appendage in less than a month (Akimenko et al., 2003; Pfefferli and Jaźwińska, 2015; Sehring and Weidinger, 2020; Tornini and Poss, 2014). Fin regeneration depends on the activation of mature cells of the stump in the vicinity of injury. Within 2 to 3 days post-amputation (dpa), a new tissue composed of undifferentiated cells, called a blastema, emerges distally to the injury site (Fig. 1C, C’). In fact, each amputated ray develops its own blastema that connects to neighboring ones to form a regenerative outgrowth. Each principal ray can thus be autonomously regrown (Marí-Beffa and Murciano, 2010; Murciano et al., 2018; Murciano et al., 2007; Suniaga et al., 2018). After three weeks already, the regenerate recapitulates most of the original appendage, approaching termination of regeneration (Fig. 1D, D’). Although the shape of the appendage is reproduced during regeneration, the new ray branching sites do not recapitulate the original positions but are shifted distally (Fig. 1D, D’) (Azevedo et al., 2011). Ray bifurcation is thus an attractive morphometric criteria to study how the biomechanical properties of a locomotory appendage are refined by external mechanical forces.

Ray growth and patterning occurs at the distal margin of the regenerate. The early dynamics of this process can be visualized by using transgenic fish and immunofluorescence analysis. Specifically, a part of the wound epithelium that is immediately adjacent to the tips of regenerating bones transcribes the *sonic hedgehog (shh)* gene (Laforest et al., 1998), which is required for ray bifurcation (Armstrong et al., 2017). This expression can be detected by imaging of a transgenic fish strain, called *shh:GFP*, which expresses green fluorescent protein (GFP) under the control of the regulatory element of *shh* (Hadzhiev et al., 2007; Zhang et al., 2012) (Fig. 1E, G). The ray bifurcation process is initiated at 3 days post-amputation (dpa) at the level of the *shh* signal (Fig. 1E, F), and branched bones can be observed at 7 dpa (Fig. 1G, H). Beside genetic regulation, ray bifurcation is influenced by mechanical factors. Specifically, ray bifurcations can also be completely suppressed by physical separation of adjacent rays during regeneration (Marí-Beffa and Murciano, 2010). Moreover, the ventral-most principal ray and the dorsal-most principal ray, which together frame the main fin paddle, have never been observed to bifurcate neither in the zebrafish nor in any other teleost fishes (Schindelin et al., 2012) (Fig. 1A, B). Transplantation experiments have highlighted the role of the interray tissue in the bifurcation process (Mari-Beffa et al., 1996; Marí-Beffa et al., 1999; Murciano et al., 2002). It is hypothesized that external mechanical forces are being transduced by the interray tissue to the underlying mechanosensing cellular components of the rays (Aiello et al., 2017), hence modulating the biomechanical structure of the growing locomotory appendage. In mammals, shear forces can be sensed through cellular mechanotransduction by osteocytes (Noda, 2011). Zebrafish osteoblasts have been suggested to have a similar role in mechanotransduction (Suniaga et al., 2018). It is not yet understood how different genetic and environmental inputs are integrated to guide the ray pattern during regeneration in presence of hydrodynamic forces.

Throughout ontogeny and regeneration, the zebrafish experiences an important transition in flow regime (McHenry and Lauder, 2006; Müller et al., 2000). The flow regime of a flapping fin can be defined based on its Reynolds number (*R = uL* / *ν*), which is defined by the average external flow speed (*u* [m/s]), the fin characteristic length (*L*, [m]) and the kinematic viscosity of the fluid (*ν* [m^2^/s]). Larvae and juvenile zebrafish operate mostly in the viscous regime (*R* <300), where viscous shear forces acting tangentially to the surface dominate (McHenry and Lauder, 2006; Müller et al., 2000). Ontogenic growth induces the transition of zebrafish through the intermediate regime (300 < *R* <1000) and into the inertial one (*R* >1000) where forces perpendicular to the fin dominate. They are at least one order of magnitude larger than viscous shear forces and they offer the main contribution to propulsive forward forces (thrust). During regeneration, the gradual expansion of the fin surface with supportive rays implies related hydrodynamic changes, where the ratio of inertial versus viscous forces increases progressively, as the length (and thus *R*) of the appendage increases. Mechanotransduction of these different forces and their respective contribution could be cues involved in regulating ray bifurcation. In particular, changes in mechanical conditions during ontogeny and regenerative growth could explain the morphological differences observed between the branching patterns in uninjured caudal fins and regenerated ones.

Measurements of spatially and temporally resolved hydrodynamic forces can be obtained by combining surface reconstructions of the analyzed object with high-resolution velocity maps of the surrounding flow of water (Fig. 2) (Dagenais and Aegerter, 2021). Flow fields can be quantified by tracking at very high resolution tracing particles seeded into the liquid, an experimental approach called particle tracking velocimetry (PTV) (Dracos, 1996; Maas et al., 1993; Pereira et al., 2006). Such reconstructions of fluid velocity fields have allowed researchers to evaluate the vortex wake and propulsive forces of synthetic foils (Bohl and Koochesfahani, 2009; David et al., 2012; David et al., 2017; Godoy-Diana et al., 2008; Green et al., 2011; Kim and Gharib, 2010; Marais et al., 2012; Muir et al., 2017; Shinde and Arakeri, 2014; Triantafyllou et al., 2004) and of elaborate bioinspired fin replicas (Dagenais and Aegerter, 2020; Dewey et al., 2012; Esposito et al., 2012; Ren et al., 2016a; Ren et al., 2016b; Tangorra et al., 2007; Tangorra et al., 2010). Hence, properly tuned biomimetic hydrofoils can provide precious information about the external mechanical forces acting on a caudal fin.

**Figure 2.**
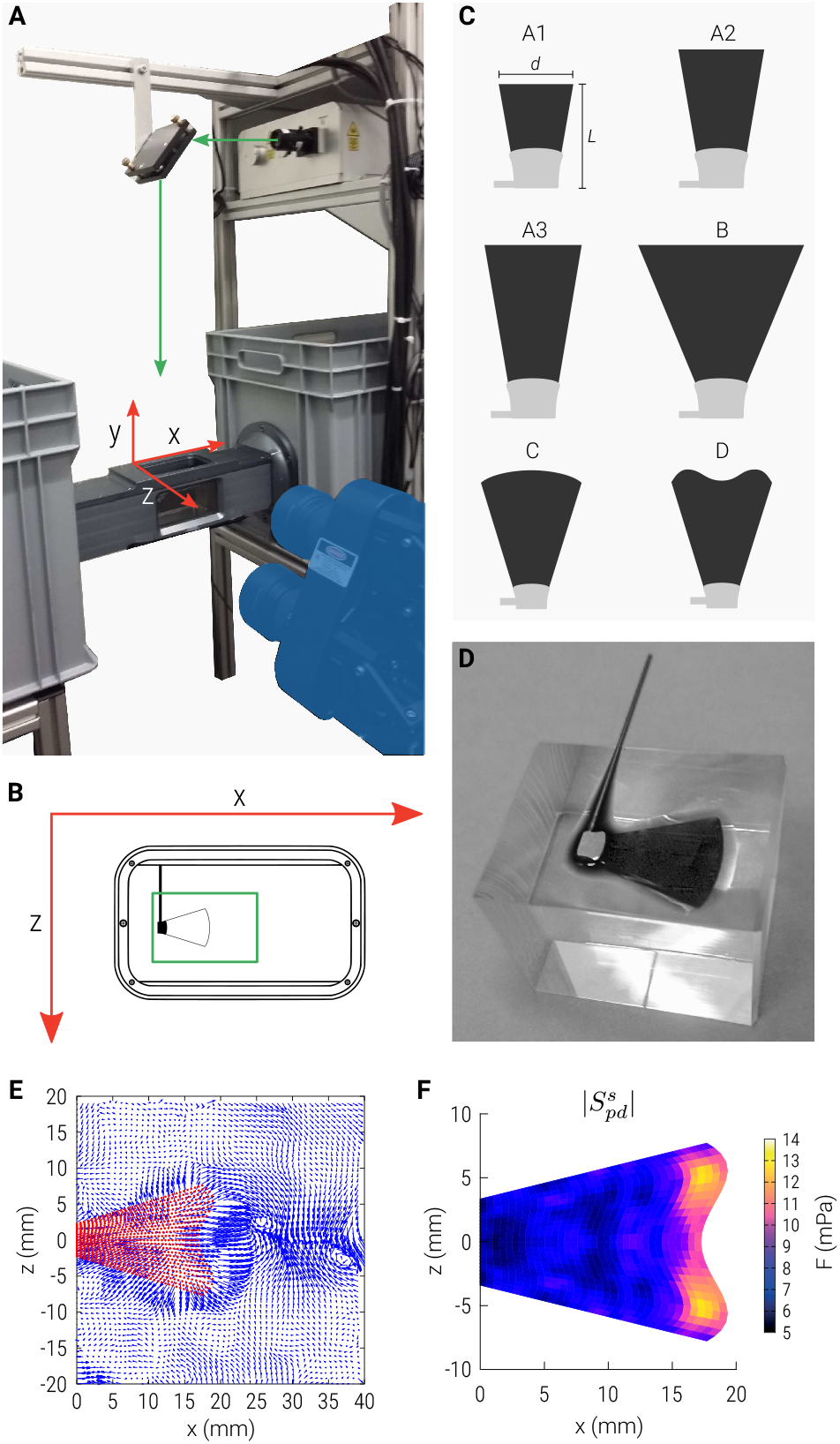
Experimental set-up with synthetic fin models for flow velocimetry experiments. (**A**) PTV set-up: laser beam (indicated by a green arrow) expanded with a pair of cylindrical lenses, deflected by a mirror to illuminate a flow chamber (x,y,z-axes indicated in red) imaged by a triplet of cameras (highlighted in blue) and connected to two water tanks (gray). (**B**) Top view of the flow chamber with measurement volume (illuminated by the expanded laser beam) indicated in green with a hydrofoil mounted on its actuating rod. (**C**) Morphology of the six fin models and their characteristic dimensions (*L* length, *d* width, Table 1). (**D**) Picture of the hydrofoil C during fabrication, as the PDMS is poured into its mold. (**E**) Fluid velocity vectors in the midplane (extracted from the 3D field), as the foil D crosses the zero-angle position, measured in the reference frame of the fin (i.e. upstream velocity subtracted from the velocity field measured in the lab frame). (**F**) Stress map of the absolute proximo-distal component acting on the side of the fin 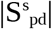. Stress intensity is color-coded according to the depicted colorbar.

In the present work, we combined surgical manipulation with caudal fin regeneration in zebrafish to modulate the biomechanical properties of the locomotory appendage. Our results indicate that the bifurcations of the bony rays are modulated by the surface area of the paddle. In addition, we performed PTV on a range of hydrofoils to quantify the distributions of mechanical forces acting at different stages of regeneration. By correlating our biological and physical data, we conclude that viscous shear stress acting on the tip of the fin and the resulting internal tension may contribute to the regulation of ray growth dynamics and branching pattern. Our combined findings reinforce the role of mechanical forces as environmental factors that modulate genetically encoded information to produce a locomotory appendage.

## METHODS

### Animal procedures and live imaging

The following strains of zebrafish *(Danio rerio)* were used in this study: wild type AB (Oregon strain) and *Tg(−2.2shha:gfp:ABC)* (Ertzer et al., 2007), referred to as *shh:GFP; (OlSp7:mCherry^zf131^)* (Spoorendonk et al., 2008), referred to as *osterix:mCherry.* Animals were maintained at 26.5 °C. Fin amputations were performed as described previously (Pfefferli and Jaźwińska, 2017). Briefly, the fish at 8-20 months of age were first incubated in analgesic solution, containing 5 mg/l lidocaine hydrochloride (Sigma-Aldrich) for 1 h before procedure to suppress pain perception. Then, the fish were anesthetized in 0.6 mM tricaine (MS-222 ethyl-m-aminobenzoate, Sigma-Aldrich) and transferred under a stereomicroscope for surgeries. Straight amputations of caudal fins were performed using a razor blade. After fin amputation, excisions of rays were conducted using a micro dissecting knife (RS-6220, dean knife 5” 1 mm x 7 mm blade curved, Roboz Surgical Instrument Co., Inc., Gaithersburg, Maryland, USA). To remove rays, incisions in the soft tissue surrounding specific rays were performed, and then, the rays were cut near the peduncle at the base of the fin. Different combinations of rays were ablated, as schematically illustrated (Fig. 3). The fins were subsequently allowed to regenerate. In order to prevent regrowth of the ablated rays during regeneration, the removal of rays was repeated every 3 to 4 days.

**Figure 3.**
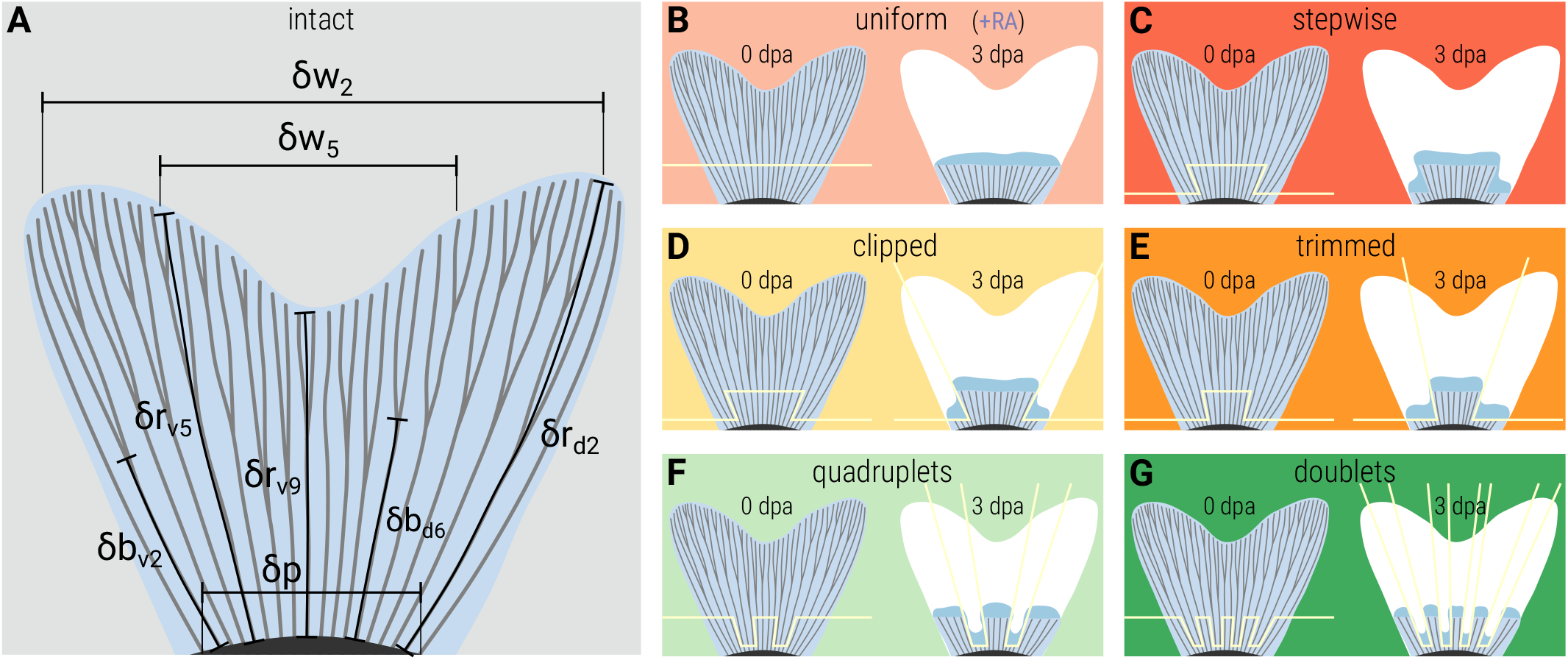
Morphometric measurements and amputation patterns. (**A**) Schematic representation of the various measurements used in this study overlaid on an uncut fin. Ray length (δr) is measured from the peduncle to the tip of the fin following the center of the area spanned by a ray (e.g. δr_v5_, δr_v9_, δr_d2_). Bifurcation position (δb) is measured similarly until the separation of the two rays (e.g. δb_v2_, δb_d6_). Interray width (δw) is measured as the distance between the tip of two rays of identical indexes (e.g. δw_2_, δw_5_). (**B-G**) Amputation strategies used in this study to challenge fin regeneration, overlaid with the corresponding naming convention. The performed amputations are depicted with yellow lines overlaid with an uncut fin at 0 dpa. **(B, C)** Fins were allowed to naturally regenerate after surgery. **(D-G)** After the amputation procedure, selected rays were ablated near the peduncle every 3 days during the duration of the experiment (21 days) to prevent their regeneration. Repeated ablations of specific rays are depicted by the yellow lines overlaying the regenerating fin at 3 dpa.

For treatment with retinoic acid (Sigma-Aldrich, St-Louise, MO, USA), we prepared a 1000-times concentrated stock solution dissolved in DMSO and stored at −20 °C. The final concentration was obtained by diluting the stock solution in fish water to achieve 1 μM retinoic acid. During the treatment, the fish were maintained in tanks without water circulation. The water was changed every second day. The fish were fed once a day with Artemia. Live imaging was performed with a Leica stereomicroscope coupled to a Leica DCF425 C camera for color images and a Leica DCF345 FX camera for fluorescence and black & white images. All animal experiments were approved by the cantonal veterinary office of Fribourg.

### Immunofluorescence analysis of whole fins

Fins were collected at different time-points after amputation and were fixed in 4% paraformaldehyde. Immunofluorescence staining of whole-mount fins was performed as previously described (König et al., 2018). The following primary antibodies were used: mouse Zns5 at 1:250 (Zebrafish International Resource Center) and chicken anti-GFP at 1:2000 (GFP-1010, Aves Labs). The secondary antibodies (at 1:500) were Alexa conjugated (Jackson Immuno Research Laboratories). Immunofluorescence was imaged using a Leica SP5 confocal microscope. Images were assembled using Adobe Photoshop.

### Morphometric analysis of the fins

Fiji was used to manually segment images of caudal fins (Schindelin et al., 2012). Average position of each bony ray was drawn using a segmented line region of interest (ROI) placed at the center of the area spanned by bifurcated rays (Fig. 3A, S1A). Position of the distal part of the ray from the peduncle arch up to the first bifurcation, or to the tip of the ray for unbifurcated rays, was drawn similarly (Fig. 3A, S1A). All segmentations were organized using the ROI manager functionality of Fiji and exported as compressed “.roi” files.

Images for the following 8 experimental groups (Fig. 3) were analyzed: intact fins (*I*, n=31, Fig. 3A, S1A), uniform amputations (*U*, n=24, Fig. 3B, S1B), retinoic acid-treated uniform amputations (*R*, n=5, Fig. 3B, 4D), stepwise amputation (*S*, n=5, Fig. 3C, S1C), clipped (*C*, n=33, Fig. 3D, S1D), trimmed *(T*, n=6, Fig. 3E, S1E), quadruplets (*Q*, n=8, Fig. 3F, S1F) and doublets (*D*, n=7, Fig. 3G, S1G).

**Figure 4.**
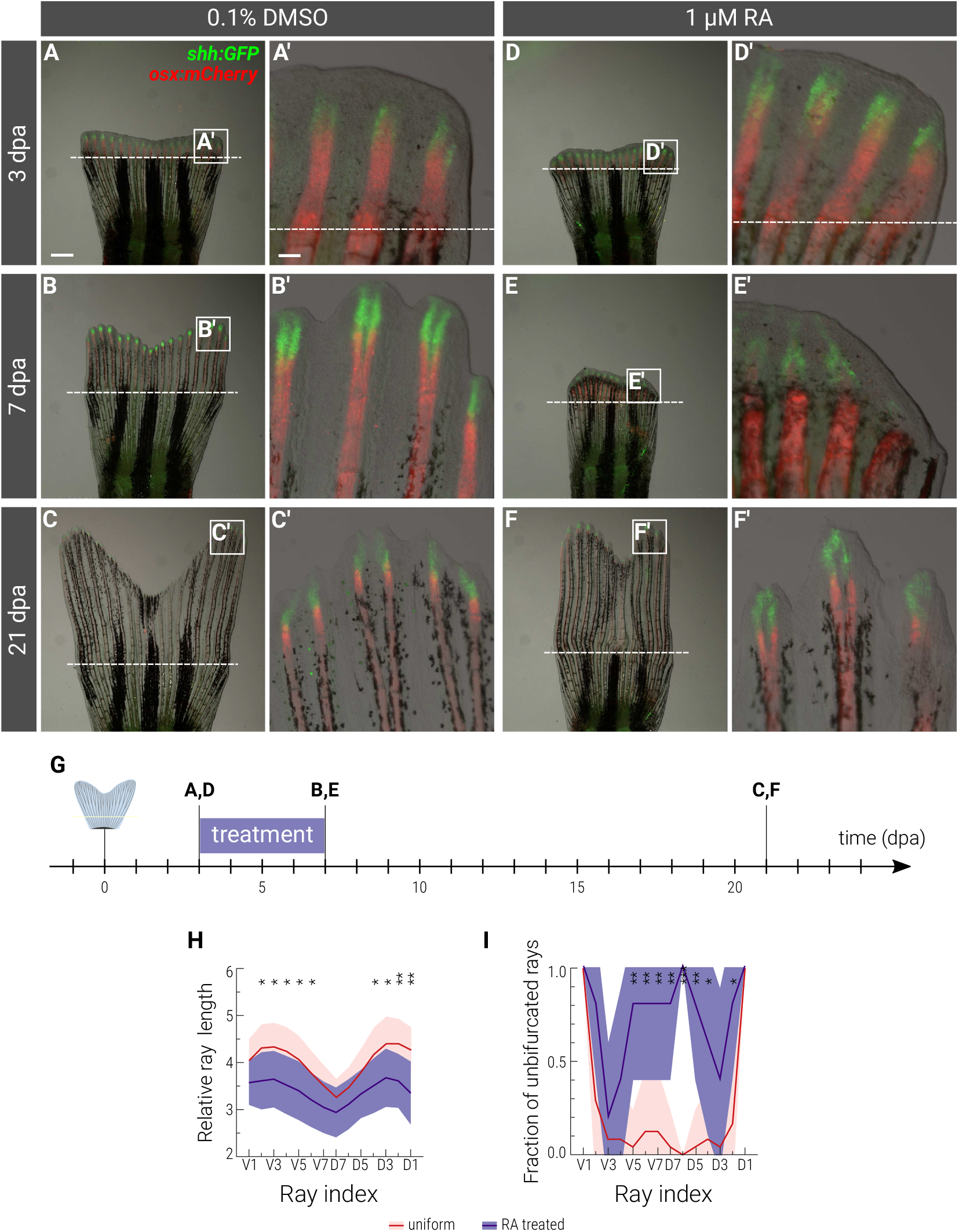
A transient retinoic acid treatment during regeneration results in narrower fins with distalized bifurcations. (**A-F**) Live-imaging of fins expressing *shh:GFP* (green); *osterix:mCherry* (red) at the indicated time points. Dashed lines depict the amputation planes. At 3 dpa, before the treatment, no difference is observed between control and manipulated groups. At 7 dpa, RA treatment blocked regeneration and caused contraction of the blastema. At 21 dpa, the regeneration is almost complete in both control and manipulated fins, although the fin that was treated with RA is much narrower than control. Scale bars is 1 mm in **A** and 100 μm in **A’**. (**G**) Experimental design with the time line post-amputation. Depicted are the amputation, the treatments and the time-points displayed in **A-F**. Control is treated with the solvent 0.1% DMSO. The manipulated group is transiently treated with 0.1 μM retinoic acid (RA) during 4 days, starting from 3 dpa until 7 dpa to contract the wound epithelium. (**H**) Distribution of ray lengths at 21 dpa, normalized to the corresponding peduncle width. (**I**) Distribution of unbifurcated rays at 21 dpa. Ray indices are provided at the bottom of the plots. Color-codes of the mean curves and standard deviations follow the legend displayed at the bottom of the figure. Significance is measured with respect to the distribution in uniform amputations using a two-tailed Student’s *t*-test for **H** and using a Fisher’s exact test for **I**. Significance is depicted as * *p*-value < 0.05, ** *p*-value < 0.01, *** *p*-value < 0.001.

Data were analyzed using the GNU Octave programming language with custom scripts. ROIs were loaded using the ReadImageJROI function developed by Dylan Muir (Muir and Kampa, 2015) and lengths measured as the Euclidian distance between the vertices of the ROIs. To compensate for the variability in ray number present in zebrafish caudal fins, only the seven outermost dorsal and ventral rays were taken into account for ray-level analyzes.

Morphometric properties were calculated, for fin measurements, normalized to the fish peduncle’s width to compensate for any variation in animal size, and for bifurcation measurements, normalized to the corresponding ray length. All relative measures were then divided by the average value in intact fins. Relative fin length: 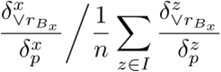, where 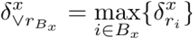, 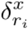 is the length of ray *i* of the regenerated fin *x, B_x_* is the set of non-ablated bony rays of fin *x*, 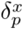 the peduncle width of fin *x* measured as the distance between the proximal vertices of the two outermost non-ablated rays, *I* the set of intact fins described above and *n* the number of fins in this experimental group (*n*=31); relative cleft depth: 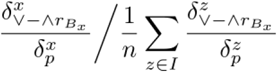, where 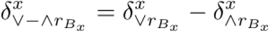 and 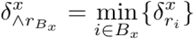; relative fin width: 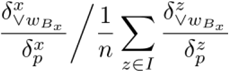, where 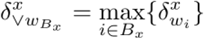 and 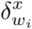 is the distance between the distal ends of dorsal and ventral rays *i*; relative bifurcation position: 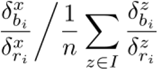, where 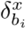 is the length of ray *i* until its first bifurcation point; normalized ray length: 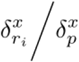; fraction of unbifurcated rays: 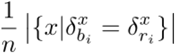.

Statistical significance was calculated using Student’s two-tailed *t*-tests for real values, and using a Fisher’s exact test for binary values. Significance is represented as * *p*-value < 0.05, ** *p*-value < 0.01, *** *p*-value < 0.001.

### Quantification of flow velocimetry

We used a three-dimensional, three components particle tracking velocimetry approach (3D-3C PTV) to quantify flow in a custom made recirculating system composed of two large tanks connected to both sides of a 8.5 × 8.5 × 12 cm^3^ flow chamber (Fig. 2A). A pump pushed water from the upstream tank into the flow chamber through a flow straightener to obtain a laminar stream of controlled fluid velocity. 3D fluid velocity vector fields were recorded as previously described (Lai et al., 2008; Pereira et al., 2006; Pothos et al., 2009). Briefly, ~50 μm polyamides tracer particles were seeded in the flow and illuminated by a 200 mJ dual-head pulsed Nd:YAG laser equipped with a pair of cylindrical lenses to expand the beam. Flow was imaged using the V3V-9800 system from TSI Inc. (Lai et al., 2008) at a temporal resolution of δt = 2.5 ms. The cameras sensors size was 11.3 × 11.3 mm^**2**^, the magnification 0.3X, the resolution 2048 × 2048 pixels and the distance to the center of the chamber ~46.5 cm. This set-up enabled us to monitor a volume of 50 × 50 × 20 mm^3^ (Fig. 2B).

Reconstruction of the particles’ positions and velocities was performed using the V3V software (Lai et al., 2008; Pothos et al., 2009). A 2D Gaussian fit of the particles intensity distributions in each image was used to identify the 2D positions maps. Triplets of position fields were captured simultaneously by the three cameras and superimposed to compute the 3D positions by triangulation. Our temporal resolution resulted in an average displacement of about 10 pixels between subsequent time frames. 3D fluid velocity vector fields were then calculated between two consecutive position fields using a particle tracking algorithm based on a relaxation method (Lai et al., 2008; Pothos et al., 2009). Raw velocity fields were filtered to remove outliers using three different filters: a velocity range filter (where vectors whose components exceed ±0.25 m/s are removed), a Gaussian smoothing filter (σ=1.5 voxels) and a spatial median filter (with a neighborhood size equal to 5 mm). Filtered 3D velocity vectors were then interpolated onto a regular Cartesian grid with an isotropic spatial resolution of 0.75 mm (Fig. 2E).

### Biomimetic hydrofoils

Six different hydrofoils designs (Fig. 2C) were produced as follows. A rigid hollow cast was 3D printed, liquid polydimethylsiloxane (PDMS) was poured inside the cast, a fan-shaped rod was inserted at the base of the hydrofoil and the material was cured for 36 hours at 58 °C. The resulting flexible membranes (Fig. 2D) have a thickness (w) of 0.55 mm and a Young’s modulus of approximately 0.8 MPa (Puri et al., 2018). The hydrofoils were used independently in the flow chamber. They were mounted horizontally inside the chamber and actuated through a dedicated opening on the back wall by a servomotor fixed outside the chamber. The flapping movement was a pitching sinusoidal motion with its rotation axis parallel to the z axis and perpendicular to the cameras plate. The exact position of the foil was determined by image analysis. In each time frame, 25 points were manually drawn on the midline of the fin, and fitted with a quadratic function. The 3D surfaces of the hydrofoil were then reconstructed from this fitted midline. The reconstructed fin surface is at most 0.7 mm away from the real surface owing to the fact that particles located at the fluid-solid interface cannot be resolved. This distance is smaller than the spatial resolution, allowing the shear stress patterns to be properly captured at the surface.

We use two dimensionless parameters to determine the flow regime acting on our flapping fin-like hydrofoils and to encompass the propulsion dependence on oscillations: *R* and the Strouhal number (*St=fA*/*u)*, where *f* is the flapping frequency and *A* the amplitude of the oscillations. More theoretical background about these parameters and the characterization of flow regimes can be found in (Prandtl, 1952; Whitaker, 1968). Given that *v* = 10^-6^ *m^2^/s* for water at room temperature, we tuned the values of the other parameters *(f=3* Hz, *u*=0.055*m/s*) and the characteristics of the hydrofoils (Fig. 2C, Table 1, 0.015≤L≤0.025 m, *A* inferred by image analysis) to fit biologically relevant values of *R* and *St* when compared to those measured for zebrafish caudal fins (Mwaffo et al., 2017; Palstra et al., 2010; Parichy et al., 2009; Taylor et al., 2003; Triantafyllou et al., 2000).

**Table 1.**
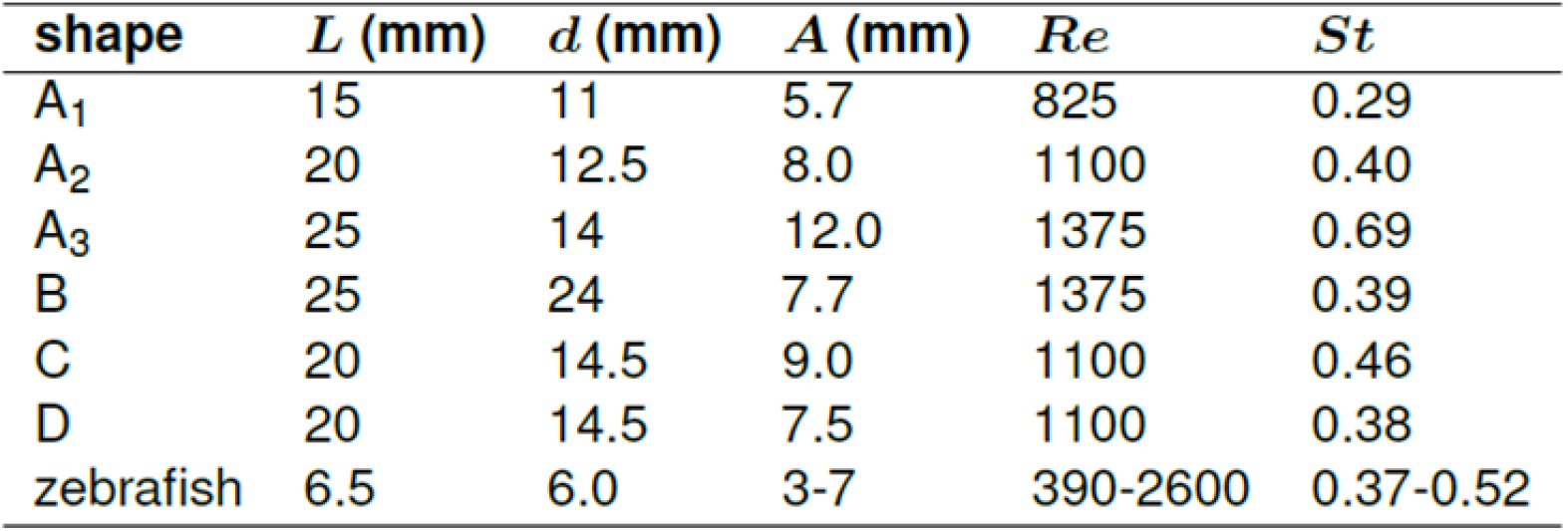
Morphometric and flow parameters of the hydrofoils and of the zebrafish caudal fin

| shape | $L$ (mm) | $d$ (mm) | $A$ (mm) | $Re$ | $St$ |
| --- | --- | --- | --- | --- | --- |
| $A_1$ | 15 | 11 | 5.7 | 825 | 0.29 |
| $A_2$ | 20 | 12.5 | 8.0 | 1100 | 0.40 |
| $A_3$ | 25 | 14 | 12.0 | 1375 | 0.69 |
| B | 25 | 24 | 7.7 | 1375 | 0.39 |
| C | 20 | 14.5 | 9.0 | 1100 | 0.46 |
| D | 20 | 14.5 | 7.5 | 1100 | 0.38 |
| zebrafish | 6.5 | 6.0 | 3-7 | 390-2600 | 0.37-0.52 |

### Surface stress maps

Time-resolved stress maps on the surfaces of flexible fins were obtained as recently described (Dagenais and Aegerter, 2021). Briefly, the viscous stress tensor at time *t* is a function of the spatial derivatives of the velocity field: 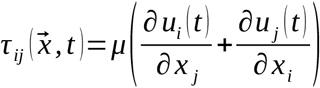. This tensor was evaluated on each vertex of the 3D cartesian grid using a finite central difference scheme. *τ_ij_* was then projected on the surface of the solid fin to obtain a stress vector, which is the total force per unit area generated by the fluid on the foil. Uniformly spaced unit vectors 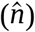 normal to the surface of the foil, and pointing outwards, were calculated based on the vertices of the 3D reconstructed fin, which consists of 25 nodes in the dorso-ventral direction, 25 nodes in the anterior-posterior direction and 3 nodes in the left-right direction. A coordinate system local to each node was defined as *(dv, ap, lr)*, where *dv* is the local dorso-ventral axis, which is coplanar with the foil’s side surface and parallel to the base-tip midline, *ap* is the anterior-posterior axis, which is perpendicular to *dv* and also coplanar with the side surface, and *lr* is the left-right axis, which is normal to the side surface. Each component of the stress vector acting on a surface with a unit normal vector 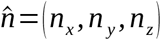 (oriented outwards) is expressed according to 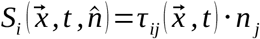. The cartesian stress vectors (*S_x_*, *S_y_, S_z_*) were finally decomposed into biologically relevant components, by projecting them onto the (*dv*, *ap*, *lr*) directions, which gives a total of six different stress components. These are the three components on the side of the foil in the local coordinate system, i.e. 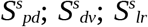 as well as on the outer tip surface of the foil, i.e. 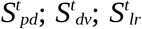. The components on the dorsal and ventral outer surfaces were not considered in this analysis because of their negligible involvement during morphogenesis.

These stress maps were evaluated at 30 time points equally distributed over 3 flapping periods, and the results were time-averaged over these 3 periods. Spatial averaging was also performed following the dorso-ventral symmetry of the hydrofoil to reduce experimental uncertainties. This averaging procedure was performed for both the absolute values and the signed values of the stresses, yielding a total of twelve different stress measures to evaluate. Signed values provide a measure of the resulting internal tension (i.e. of the excess of contraction or stretching at the surface). Due to symmetries in the forces acting on the fin during a flapping period as well as the symmetric shape, spatial averaging yields to 0 for the stresses 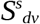 and 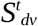. To determine the tension in the *dorso-ventral* direction, we calculated 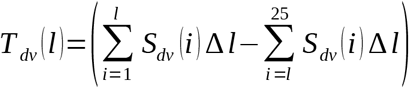, where *l* is the position along the *dv*-axis and Δ*l* the spatial step size between nodes on the surface of the foil. With this, positive values of 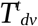 and 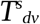 (with units [*μN/m*]) indicate contraction whereas negative values indicate stretching. 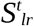 has a period average equal to zero, due to the flapping motion which is symmetrical in the left-right direction. This component is therefore left out of the analysis. Hence, 11 stress components were precisely quantified: 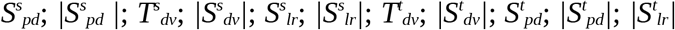. To refine our analysis of stress distributions, and using the proportions of uninjured fins (Fig. 1A), we classified the 25 nodes of our stress maps along the *dv*-axis as follows: 2 nodes for the dorsal rim, 8 for the dorsal lobe, 5 for the cleft, 8 for the ventral lobe, 2 for the ventral rim. Using this definition, we can formalize averages of the stress components in different regions for the screening as follows: 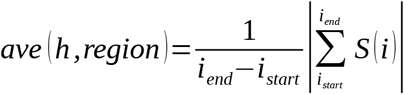, where *h* is the hydrofoil, which *S(i)* is measured on, *region* refers to the lobe, cleft or rim and *i_end* and *i_start* are the beginning and end points of the above defined regions. For visualization purposes, averages were normalized to the corresponding values on hydrofoil A2.

A detailed analysis of experimental uncertainties and limitations in resolution for our experimental set-up was previously published (Dagenais and Aegerter, 2021). Briefly, error analysis of three-dimensional velocimetry systems (Graff and Gharib, 2008) was used to infer the precision of our measurements. Prior to performing the particle tracking experiments, the system was calibrated to derive the cameras model by using a plate with a known grid pattern, imaged by the three cameras at different locations so that the entire volume was scanned. Positional uncertainties inherent to PTV are a function of particle size, magnification, 2D Gaussian fit during particle identification, least squares error (LSE) from the calibration and number of cameras. Using the sum of squares error propagation method, the positional uncertainties were calculated by the manufacturer for all directions (x, y, z) to be (3.6, 3.6, 32) μm. Based on these positional uncertainties, and assuming that the time difference δt is exact, we obtained the error on velocity components: *σ*=(2, 2, 18) mm/s. The errors were reduced through temporal and spatial averaging and then propagated to the corresponding measures. These uncertainties are displayed as error bars in all our graphs.

## RESULTS

### Modulating the biomechanical properties of regenerated fins

Fin regeneration recapitulates the shape and size, but is associated with a distal shift in the position of the bifurcations of the regenerated rays relative to those in the original appendage (Azevedo et al., 2011). We consistently observed that a straight amputation of the caudal fin, here termed uniform amputation (Fig. 3B, S1B), induces a distal shift (Fig. 1). We hypothesized that this displacement is a consequence of an alteration of the hydrodynamic forces acting on the forming tissue compared to those present during ontogenesis. To test this hypothesis, we set out to modify the biomechanical properties of the stump and assess whether it alters the regenerated fin and the bony rays. Indeed, geometrical changes are expected to alter the hydrodynamic forces acting on the surfaces of the appendage, including the region of forming tissue (Dagenais and Aegerter, 2021; Feilich and Lauder, 2015; Low and Chong, 2010; Lucas et al., 2017; Matta et al., 2020; Valdivia y Alvarado and Youcef-Toumi, 2005).

Surface area of the caudal fin is the most readily manipulable property of the caudal fin. We devised two complementary approaches to obtain narrower fins. The first one is a surgical procedure, termed stepwise amputation, where a uniform amputation is followed by a more proximal amputation of the three dorsal and three ventral outermost rays (Fig. 3C, S1C). The second one is a non-invasive transient treatment of the blastema with retinoic acid (RA, Fig. 4), which has been shown to deform the regenerating fin by contracting the wound epidermis (Ferretti and Geraudie, 1995; White et al., 1994). For this experiment, we used double transgenic fish *shh:GFP*, demarcating the signaling center of the wound epithelium, and *osterix:mCherry*, highlighting activated osteoblasts (Knopf et al., 2011). Following a pulse treatment with chemicals, fins of both groups were allowed to regenerate in a natural way for 21 days (Fig. 4A-G).

Consistent with previous studies (Lee et al., 2005; Monteiro et al., 2014; Uemoto et al., 2020), we observed that more proximally amputated rays regenerated at an enhanced speed in comparison to distally cut rays (Fig. S1C). At 21 dpa, morphometric analysis of fin regenerates indicated that the fins recreated more than 80% of the original intact length, being close to the termination of regeneration (Fig. 5A). Relative fin width confirmed the validity of both our approaches as we observed a significant 20 % contraction in the stepwise group and a further 30 % contraction in the RA group (Fig. 5B). Some of the rays seemed to have fused following RA treatment (Fig. 4F), highlighting the substantial molecular as well as cellular plasticity of the blastema and confirming this approach for chemical ray ablation. Interestingly, we also measured an average 20 % shallower cleft in both groups when compared to uniform amputations (Fig. 5C). Importantly, while distalization of the bifurcations in the stepwise group was comparable to that of the uniform group, RA-treated fins displayed a significant further distalization (Fig. 5D). Overall, these results confirmed that caudal fin morphology is modulated by the manipulation of its biomechanical properties during regeneration.

**Figure 5.**
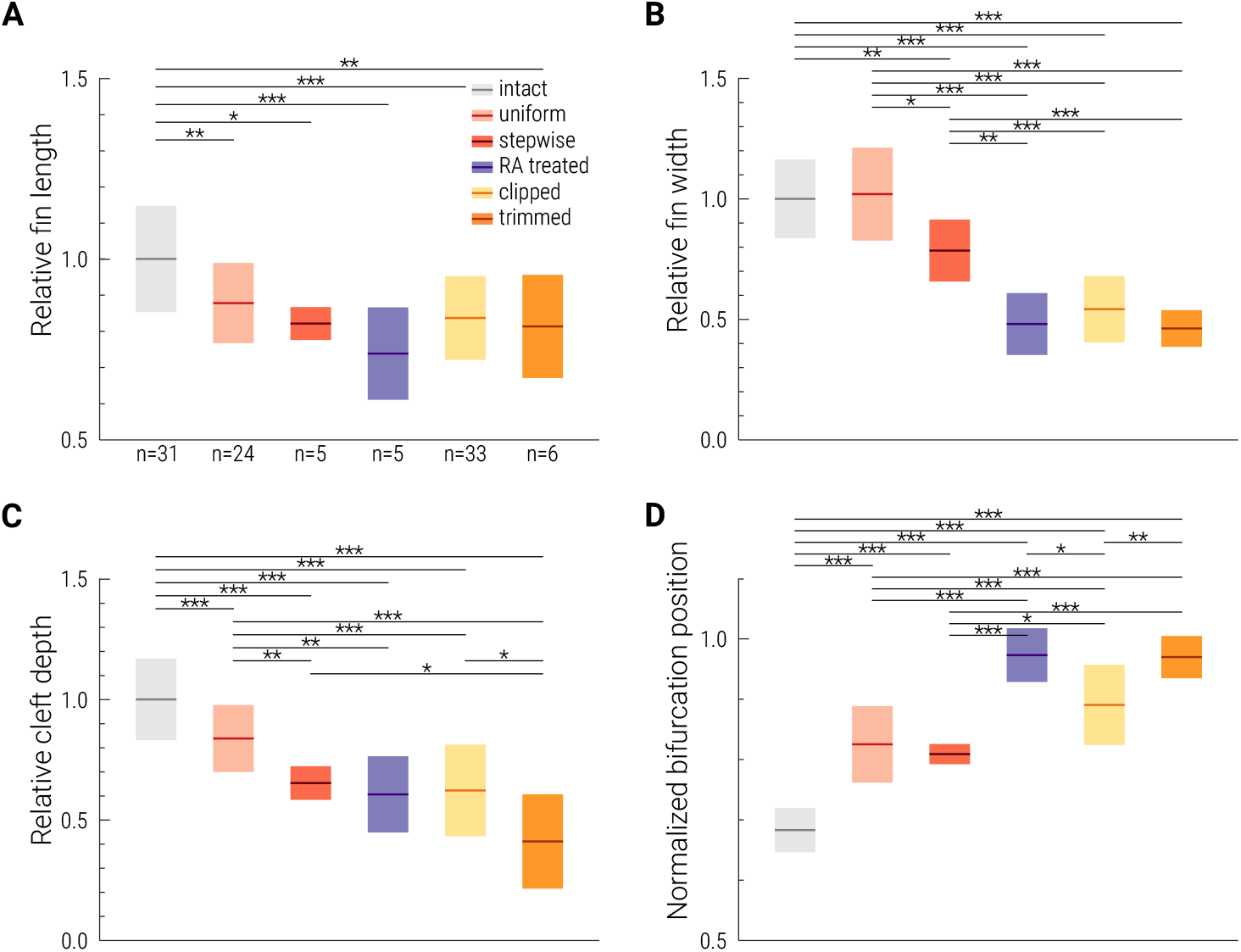
Impact of the amputation strategies on the morphological properties of regenerated fins. (**A**) Fin lengths at 21 dpa relative to the corresponding peduncle width. (**B**) Relative fin widths. All conventions follow those in **A**. (**C**) Relative cleft depths. (**D**) Bifurcation position normalized to the corresponding ray length. Number of samples in each box plot is given below each group. Color-codes follow the provided legend. Calculation of all values is described in the methods. Mean values and standard deviations are shown. Significance is measured using a two-tailed Student’s *t*-test and is depicted as * *p*-value < 0.05, ** *p*-value < 0.01, *** *p*-value < 0.001.

### Phenotypic alterations in narrowed fin regenerates

The increased distalization observed following retinoic acid (RA) treatment (Fig. 5D) suggested that it was possible to enhance the phenotypic changes induced by our stepwise amputation. Because RA regenerates are significantly narrower than stepwise amputations (Fig. 5B), we hypothesized that reduced fin surface would induce such changes. We verified this hypothesis by surgically manipulating caudal fins. We thus created artificially narrower regenerates by ablating a number of the dorsal-most and ventral-most principal rays. In the clipped experimental group, the blastema of the three most ventral and the three most dorsal rays were repeatedly amputated throughout regeneration (Fig. 3D). In the trimmed one, five rays on each side were similarly prevented from contributing to the regenerated locomotory appendage (Fig. 3E).

At 21 dpa, these fins restored their expected length, as compared to uniform and stepwise amputated fins (Fig. 5A, S1D-E). In addition, these ablations were able to further narrow the relative width of the regenerates to an extent similar to the contraction observed following RA treatment (Fig. 5B). In the clipped group, we observed a shallow cleft depth comparable to both RA treatment and stepwise amputation (Fig. 5C), but a distalization phenotype intermediate to these two groups (Fig. 5D). In the trimmed group, we detected a further reduction in cleft depth and an increased distalization with respect to clipped fins (Fig. 5C-D). Analysis of single ray lengths showed that the reduced cleft depth in the clipped group was caused by a significant reduction of lengths of the outer rays (Fig. 6A). In the trimmed group, the outer rays also seemed to contribute most the observed reduction, but the outer rays were not significantly shorter (Fig. 6A). This second reduction in surface area thus reached morphological changes similar to those observed after a pulse treatment with RA (Fig. 4H). More importantly, these results show that by using mechanical alterations in the morphological properties of the regenerating appendage, we are able to induce a range of quantitative modulations of the caudal fin.

**Figure 6.**
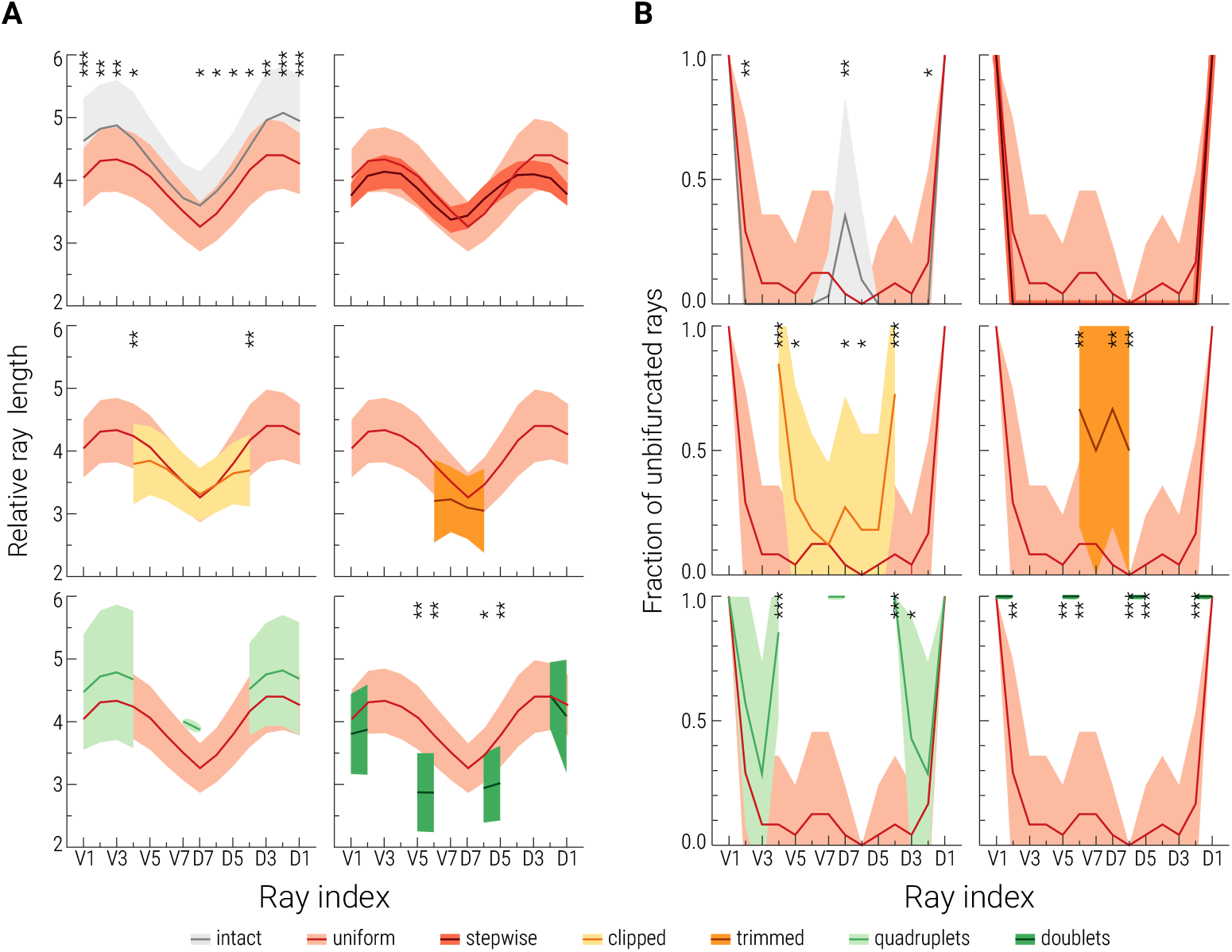
Impact of the amputation strategies on the properties of individual rays. (**A**) Distribution of ray lengths at 21 dpa, normalized to the corresponding peduncle width. (**B**) Distribution of unbifurcated rays at 21 dpa. Ray indexes are provided at the bottom of the plots. Color-codes of the mean curves and standard deviations follow the legend displayed at the bottom of the figure. Significance is measured with respect to the distribution in uniform amputations using a two-tailed Student’s *t*-test for **A** and using a Fisher’s exact test for **B**. Significance is depicted as * *p*-value < 0.05, ** *p*-value < 0.01, *** *p*-value < 0.001.

To explore how far phenotypic variations can be affected, we reduced further the width of the regenerates. We thus separated the rays of the stump by quadruplets and doublets (Fig. 3F, G). Following uniform amputation, intermediate rays were regularly ablated to prevent their regeneration (Fig. S1F, G). In addition to providing biological replicates within the same fin, this tuplet approach allowed us to investigate the influence of interray tissue on ray bifurcation (Fig. 7). Comparison of relative ray length at 21 dpa to uniform amputation confirmed that all bony rays in the tuplet experiments regenerated to their expected length (Fig. 6A), with the exception of the two central doublets. We observed an apparently random uneven length between various pairs of these central doublets, likely a result of spontaneous breakage of one of the regenerates due to the fragility of these thin structures. To refine our analysis on bifurcation distalization, we quantified bifurcation events per individual ray (Fig. 6B, Fig. 7G-I’). The fraction of unbifurcated rays per experimental group supports previous reports that the outermost principal rays do not bifurcate (Mari-Beffa et al., 1996; Marí-Beffa et al., 1999; Schultze and Arratia, 2013). Inhibition of bifurcation was confirmed at the molecular level in transgenic fish *shh:GFP*, which normally display a split *shh*-expression pattern proceeding ray splitting (Fig. 1E-H). We found that the *shh*-expression domain of the new unbifurcated outer rays surrounding the ablated regions didn’t split, again highlighting the substantial phenotypic plasticity of the blastema upon altering circumstances (Fig. 7A-F). However, our tuplets experiments also showed that while these new outer rays rarely bifurcate, they occasionally do so (Fig. 6B, 7H). These results suggest that mechanotransduction of external forces through the interray tissue contributes to splitting of *shh*-expression and that the presence of interray on both sides of a ray is not strictly required for the induction of ray branching events.

**Figure 7.**
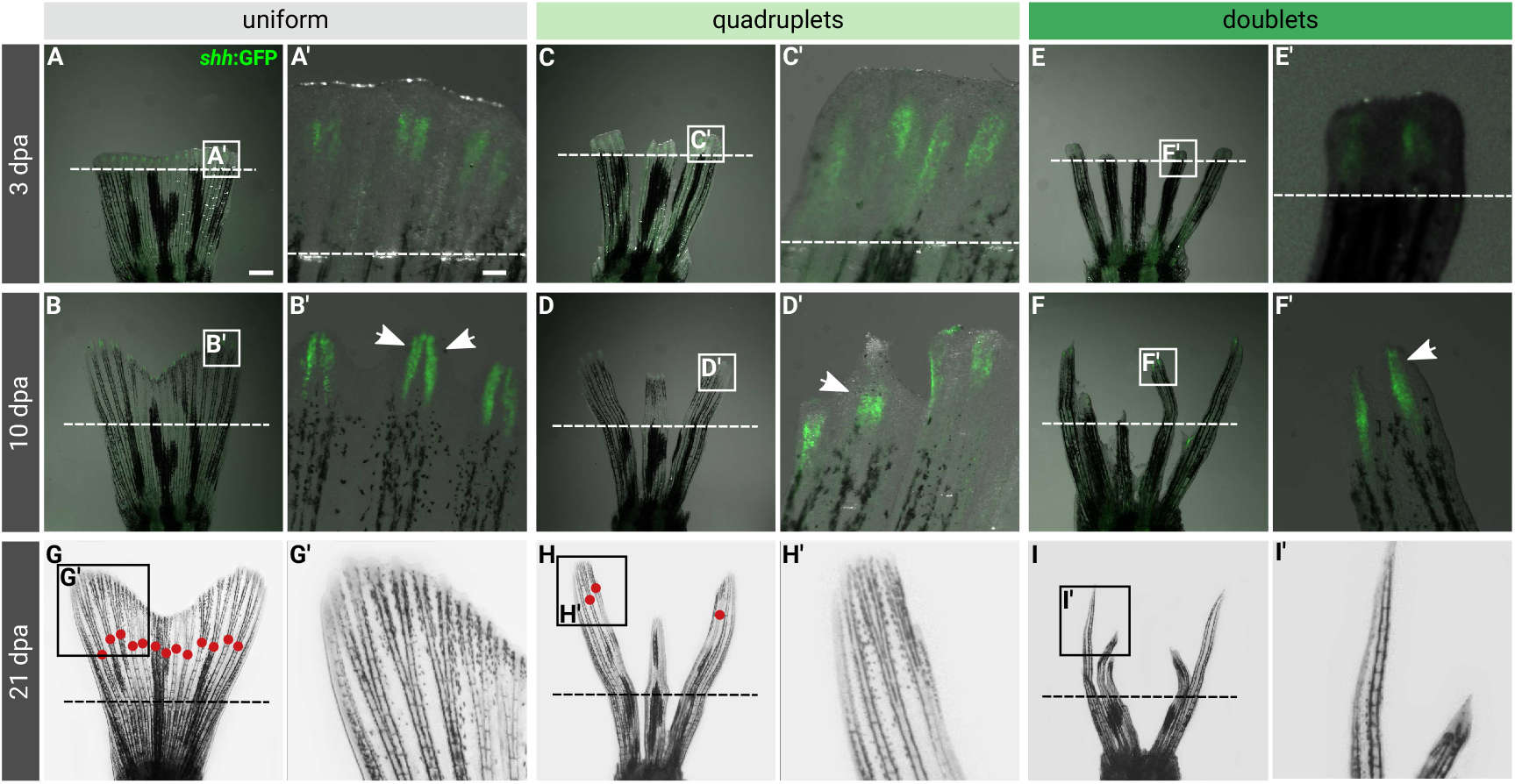
Surgical transformation of fins into ray-doublets and ray-quadruplets during regeneration suppresses bifurcation. (**A-F**) Overlaid fluorescence and bright-field images of regenerating transgenic *shh:GFP* (green) fins at the indicated time points, with the indicated cutting pattern and the amputation plane depicted by a dashed line. (**A’-F’**) Higher magnifications of the area highlighted in the corresponding images by a white rectangle. Split *shh:GFP*-expression regions are indicated by two arrows, whereas a uniform *shh:GFP* region is pointed by a single arrow. (**G-I**) Bright-field images of the manipulated fins at 21 dpa. Bifurcations are depicted with red dots. The amputation plane is shown with a dashed line.

Overall, our analysis highlights the phenotypic plasticity of the caudal fin upon biomechanical manipulation, potentially to adapt the locomotory capacity of the growing appendage.

### Investigating external hydrodynamic forces using biomimetic hydrofoils

In order to identify a potential source for mechanical feedback, we investigated whether the distributions of hydrodynamic stresses were consistent with the measured morphological modulations. To increase the precision and reproducibility of our physical measurements, we designed a set of biomimetic hydrofoils that recapitulate the most relevant features of our experiments on caudal fin regeneration (Fig. 2C, Table 1). Three foils with increasing lengths (60 %, 80 % and 100 %) were used to study how forces vary during morphogenesis. Foil A1, the shortest of these foils, replicates the fin geometry right after uniform amputation by following the proportions of a zebrafish caudal from its peduncle to the amputation plane. Foil A2, the intermediate length, corresponds to the proportions at the level of ray bifurcation. Foil A3, the longest one, represents a fully regenerated fin. To incorporate our observations with altered fin width, we designed foil B to be 70% wider than our reference foil A3. Finally, foils C and D were created to assess the influence of the cleft on the distribution of hydrodynamic forces, in particular when the distal edge undergoes a transition from flat, to slightly rounded and to bilobed during regeneration. We quantified the full hydrodynamic stress tensor acting on the surface of these differently shaped hydrofoils by combining PTV with shape reconstruction of the flapping foil within a flow chamber (Fig. 2A). This allowed us to quantify six relevant projections of stresses onto the surfaces of the foil: three for the lateral side and three for the tip. Taking the average over a flapping period of the absolute value as well as the signed values of these stresses gives a total of eleven possible hydrodynamic signals that could be involved in mechanical control of the positioning of bifurcation and growth (see methods for a detailed description). Using the different lengths (A1, A2, A3), widths (A3, B) and edge shapes (B, C, D) we can correlate changes in the hydrodynamic signals with experimental observations of growth and bifurcation position (Fig. 5, 6).

To eliminate stress components that cannot be associated with mechanical control of growth, we first considered that stress components regulating growth rates should reproduce a highly stereotypic features of fin morphogenesis: higher growth rates in shorter fins (i.e. enhanced regeneration speed in proximal cuts). This means that when comparing the average signal for the shapes A1, A2, and A3 a monotonic dependence needs to be present in order to lead to growth deceleration (increasing in the case of an inhibitor, i.e. *ave(A1, rim & lobes & cleft)<ave(A2 rim & lobes & cleft)<ave(A3 rim & lobes & cleft)*, decreasing in the case of an activator, i.e. *ave(A1 rim & lobes & cleft)>ave(A2 rim & lobes & cleft)>ave(A3 rim & lobes & cleft)).* This allowed us to classify two putative activators, seven putative inhibitors and two irregular stress components (Fig. 8A). Next, we considered the bilobed shape that the regenerating tissue acquires a few days post uniform ablation (Fig. 1C, E) and hypothesized that it results from differences in the growth patterns of the blastema. This implies that the observed shallower cleft depth in narrower fin (Fig. 5C) results from reduced growth rates in the lobe of narrower fins. Hence, the difference in the signal in the lobes vs the signal in the cleft has to change in accordance with being an inhibitor (negative) or activator (positive) and has to increase in absolute value from shape A3 to B, i.e. *|(ave(B,lobes)-ave(B, cleft))|* > *|(ave(A3,lobes)-ave(A3, cleft))|*. This highlighted three putative activators (Fig. 8B), among which only 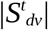 was consistent between both criteria.

**Figure 8.**
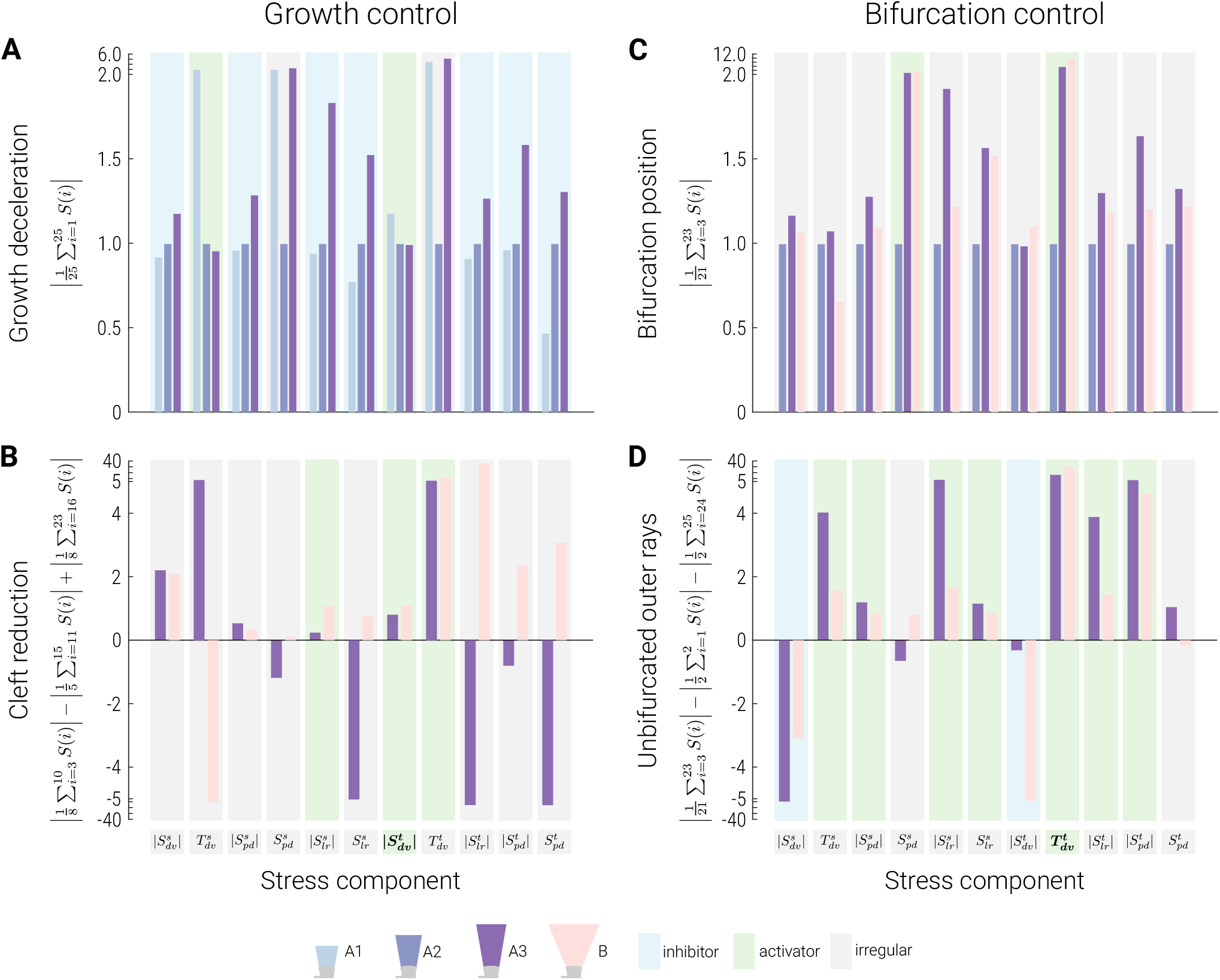
Screening stress components to identify candidate modulators of growth and bifurcation. (A) Putative controlling factors of growth need to lead to a deceleration of growth with increasing fin length from shape A1, to A2 and to A3. Activators present monotonic decreases in mean influence, inhibitors monotonic increases while other non-monotonic irregular combinations are not consistent with such a control. **(B)** Reduction in cleft depth in narrowed fins present positive and increasing values (from A3 to B) of relative lobe influence for an activator signal, while inhibitors negative and decreasing values. **(C)** For positioning of bifurcations, a signal in the lobes and cleft needs to show a consistent trend from shapes A2 to A3 and A3 to B, monotonically increasing for an activator and monotonically decreasing for an inhibitor. The comparison between A3 and B takes into account the experimentally observed distalization in narrowed fins. **(D)** To account for the nonbifurcating outer rays, the bifurcation positioning signal needs to differently influence the outer nonbifurcating rays compared to the bifurcating lobe ones. Activators show a positive signal, whereas inhibitors show a negative signal. Stress components are indicated at the bottom of panels **B** and **D**. Stress components consistent with a role in growth or bifurcation promotion are highlighted in bold. Color-codes follow the legend displayed at the bottom of the figure.

To identify a modulator of bifurcation, we are looking for a signal whose dependence on fin length is in accordance with the positioning of the blastema at 75 ± 6 % of the fin’s final length when rays bifurcate on average (Fig. 5), as well as a distalization in narrowed fins (Fig. 5D). A possible signal for this needs to have the same qualitative change when comparing the average in shapes A2 and A3, as well as shapes A3 and B. Therefore there has to be a monotonic change (increase for an activator, i.e. *ave(A2, lobes & cleft) < ave(A3, lobes & cleft) < ave(B, lobes & cleft)*, and decrease for an inhibitor, i.e. *ave(A2, lobes & cleft) > ave(A3, lobes & cleft) > ave(B, lobes & cleft))*, when comparing the averages of the stresses excluding the edges, which never bifurcate, in A2, A3, and B (Fig. 8C). This is only observed for 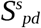 and 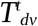. Finally, in order to be consistent with the absence of bifurcation in the outermost rays, the signal needs to differ at the edges consistently. This means that for shapes A3 and B, the difference *ave(lobes & cleft) - ave(rim)* needs to be positive in case of an activator and negative in case of an inhibitor (Fig. 8D). This leaves only 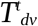 as a suitable signal, which we thus study further.

Overall, we have identified the dorso-ventral shear stress at the tip of the fin as the only one, among all hydrodynamic forces acting on the hydrofoil, displaying changes in response to length and width that are compatible with a purported mechanical feedback (Fig. 8). In particular, its absolute value 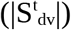 is consistent with an action as growth activator and its corresponding tension 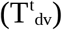 with an activation of ray bifurcations. This stress component is particularly relevant in the context of fin morphogenesis, as it corresponds to the viscous forces that can contract or stretch the tip of the growing tissue.

### Viscous shear stress distributions at the fin tip

Having identified two components of the dorso-ventral stresses at the tip of the fin as candidates for the regulation of morphological changes during growth, we studied their properties in more detail.

The first component, the absolute value of the dorso-ventral shear stress at the tip 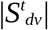, indicates which parts are subjected to a putative growth activating signal (Fig. 9A). As indicated in the screening for signal candidates (Fig. 8A), foils A1, A2 and A3 recapitulate fin regeneration. The 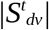 profile displays a bilobed signature in A1 already (Fig. 9A), with higher signal in the locations of the prospective lobes and lower magnitude in the center as well as on the rims. This profile offers a suitable mechanical cue for promoting growth speed as it correlates with the remodeling of the trailing edge from a straight plane to a bilobed shape, induced by variable growth rates of individual rays (Uemoto et al., 2020). Note that this consistency with an overall bilobed shape was selected in our criteria only for A3 and B, yet it is fulfilled by the signal in A1 as well as A2. As the fin model elongates (A1 to A2 to A3), 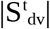 decreases in agreement with the growth deceleration observed in real fins as they approach their final size (Lee et al., 2005). The second component, 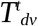 (Fig. 9B), is the putative bifurcation regulator and represents dorso-ventral contractions of the tissue when positive and conversely stretching when negative. We observed a reversal from stretching (in A1) to contraction (in A3) that correlates with the appearance of bifurcations in the blastema of fins with length similar to A2. Moreover, we measured extremely small amplitudes of tension at the location of the outermost nonbifurcating rays. Taken together, these data suggest that bifurcation is triggered by contractions of the tissue above a certain threshold.

**Figure 9.**
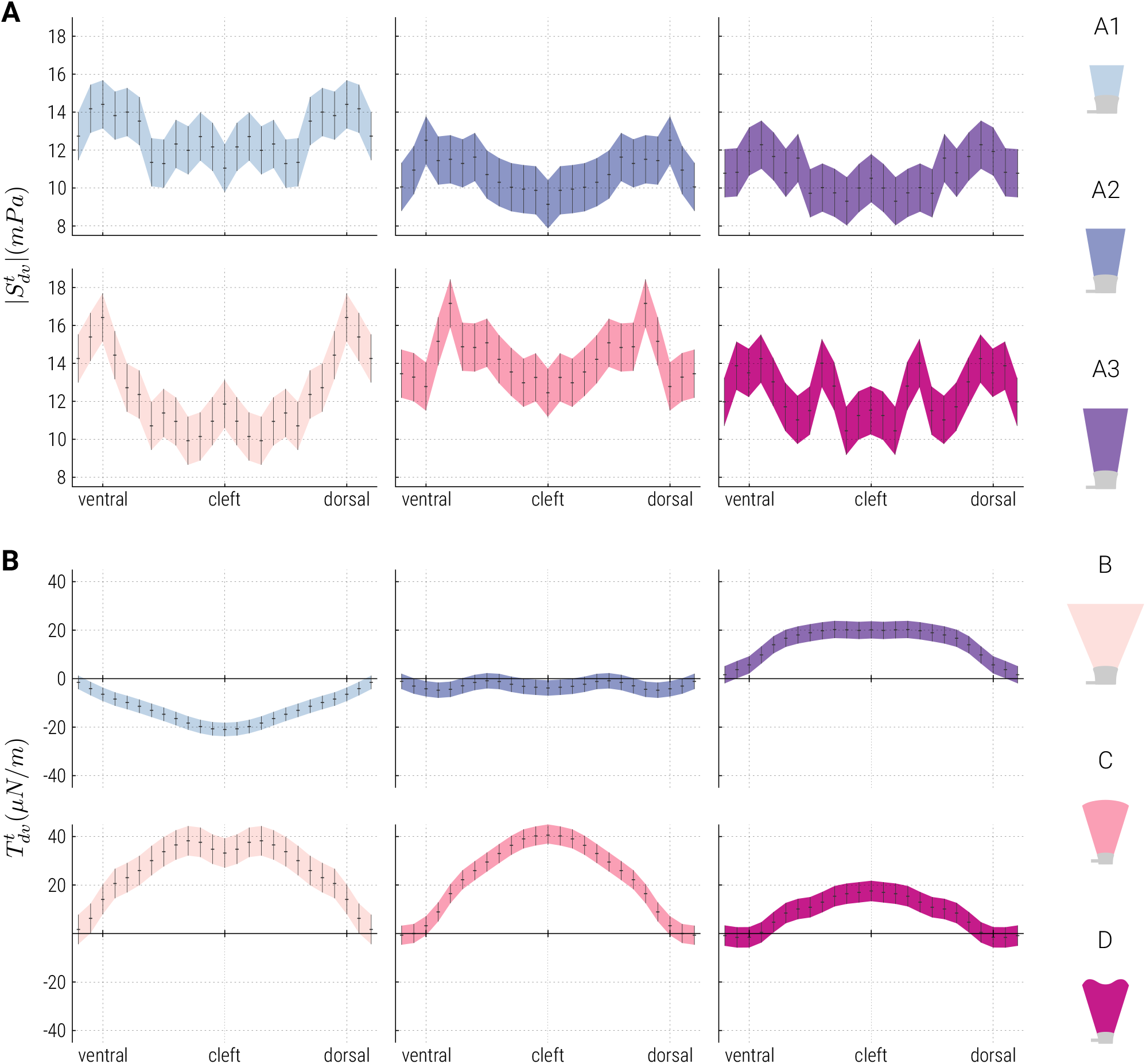
Spatial distribution of dorso-ventral shear stress at the tip of hydrofoils. (A) Absolute value of the dorso-ventral stress component 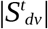. **(B)** Tension 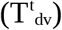. Color-codes of the mean curves and error bars corresponding to the various hydrofoils follow the legend displayed at the right of the figure.

Comparisons between a broader fin (foil B) and a narrower one (A3) highlight an increase in the magnitude of both 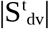 and 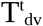 (Fig. 9). Moreover, the difference in stress magnitude between the lobes and the cleft is more important for the wider foil compared to the narrower version (Fig. 9A). Following our hypotheses, this indicates a reduction in growth rates in narrower fins, in particular in the lobes, which is consistent with our results of shorter rays in the narrower clipped fins (Fig. 6A) as well as with the measured reduction in cleft depth (Fig. 5C). Lower tension in narrower fins (Fig. 9B) is compatible with the experimentally observed distalization of bifurcation (Fig. 5D) as the length required to reach the hypothesized contraction threshold would be greater. Moreover, this lower tension would also predict the observed increase in the fraction of unbifurcated rays at the center of the fin (Fig. 6B).

Similarly, transitions from a uniform amputation (A2) to a rounded (foil C) and finally bilobed (foil D) shape show that both 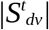 and 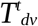 transiently increase in shape C (Fig. 9). For 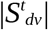 in particular, these modulations indicate a promotion of faster growth in the initial phase of regeneration, while the blastema at the tip is slightly rounded, followed by the diminution of the growth promoting signal as the final bilobed shape is attained (Fig. 9A). These variations in magnitude between the lobes and the cleft as the bilobed shape emerges correlate with the observed initial faster growth of the two lobes followed by the homogenization of growth rates towards the end of regeneration (Uemoto et al., 2020).

Overall, these findings show that the dorso-ventral shear stress component at the fin tip not only correlates with our measures of tissue growth and ray bifurcation, but that it also accounts for several more subtle experimental observations. The results thus suggest that dorso-ventral shear stress at the fin tip modulates tissue growth rates and that ray bifurcation may be triggered by tissue contraction above a certain intensity threshold.

## DISCUSSION

The caudal fin yields powerful as well as complex locomotion and is thought to be one of the most important evolutionary innovations in the aquatic environment (Flammang and Lauder, 2009). In the present work, we explored the role of mechanical forces during its morphogenesis as recapitulated by regeneration. Through a variety of surgical and chemical treatments, we have shown the phenotypic plasticity of the biomechanical structure of the fin. RA treatment induced the fusion of numerous rays (Fig. 4F), thus producing fins with a reduced number of rays. The obtained morphologies were comparable to those resulting from surgically manipulated fins with a similar number of remaining rays (Fig. 5). In particular, we have observed a distalization of the ray bifurcations and a reduction of the depth of the cleft in ablated regenerates. These changes could be adaptations aimed at improving the thrust or efficiency of the locomotory appendage by modulating the flexibility of the fin (Zhu and Bi, 2017). Such fine-tuning highlights the feedback mechanism between abiotic forces and locomotory organs. Consistently, the role of mechanical factors has also been linked with skeletal plasticity of the fish skeleton (Grünbaum et al., 2012). Several studies demonstrated that water velocity and exercise, which exert mechanical stress, influence body shape of fish, increase the number of bone-forming osteoblasts and result in higher levels of bone mineralization (Fiaz et al., 2012; Gunter and Meyer, 2014; Palstra et al., 2010; Suniaga et al., 2018). Although osteoblasts have been proposed as good candidates for the detection of mechanical loads (Witten and Hall, 2015), deciphering the mechanisms of cellular mechanotransduction, or how exactly cells transform mechanical stimuli into electric or chemical signals, constitutes a complex and unresolved problem (Suniaga et al., 2018).

Up to now, researchers have devoted most of their attention to the molecular and genetic processes related to positional information in the fin (Sehring and Weidinger, 2020; Wehner and Weidinger, 2015). Various grafting experiments have shown that position-dependent mechanisms are important for the control of the bifurcation process and the determination of individual ray lengths (Murciano et al., 2002; Murciano et al., 2007; Shibata et al., 2018). Rays grafted into a new position within the fin adopt length characteristics of the host region following regeneration, although elements of positional information also exist within the rays, independently of their host environment (Shibata et al., 2018). Another striking sign of positional information is the growth rate dependence on the proximo-distal location of the amputation plane, where more proximal structures regrow faster (Lee et al., 2005; Uemoto et al., 2020). Our study further extends these findings and indicates that biological transformations occurring during fin morphogenesis may be induced by changes in the surface distribution of hydrodynamic stresses acting on the growing blastema.

Narrower fins created by the ablation of the outermost rays undergo a shortening of the remaining lateral rays and thus present a less pronounced cleft (Fig. 5C). In these artificially thinner appendages, frequent bifurcation failure is observed in rays which normally bifurcate, most prominently in the outer rays deprived of their lateral neighboring rays, but also in the central rays (Fig. 6B). Moreover, an increased distal shift of the bifurcation points is induced (Fig. 5D). Isolating the rays by doublets or quadruplets showed similar results in terms of bifurcation failure and enhanced distal shift (Fig. 6B, 7). Previous studies have demonstrated the important role of soft tissue for bifurcation induction during fin regeneration (Mari-Beffa et al., 1996; Marí-Beffa et al., 1999). It has been shown that a single regenerating ray, deprived from any interray region, failed to undergo branching morphogenesis (Murciano et al., 2002). However, this published role of the interray tissue does not explain our observations of a further distalization of bifurcation (Fig. 5D). Consequently, we hypothesized that environmental forces could be the trigger for bifurcation and that changes in the shape and hydrodynamic conditions of the regenerate would induce the distalization.

To test this hypothesis, we measured fluid forces on the surface of hydrofoils reproducing the flow regime of caudal fins. This precise quantification demonstrated that the different hydrodynamic properties of the various fin shapes induce different force distributions. By screening for stress components compatible with the results of our regeneration experiments, we identified the dorso-ventral shear stress acting on the tip of a fin and the resulting tensile force as candidate activators of growth and ray bifurcation, respectively (Fig. 8). The results of this hydrodynamic study allowed us to put in perspective the results of our regeneration experiments on mechanically altered fins with the spatial distribution of this viscous force component. On one hand, the enhanced shear stress present at the lobes of short fins (A1, Fig. 9A) could explain the formation of a bilobed trailing edge from a flat amputation plane. Growth rate would then gradually decrease following the reduction in stress intensity as the appendage approaches its final size (A2, A3, Fig. 9A). Simultaneously, growth would reduce as it reaches its bilobed shape (A2, C, D, Fig. 9A) because the stress signal decreases predominantly in the lobes upon the transition to a rounded and to a bilobed shaped fin. The increase of the lobe signal in wide fins (A3, B, Fig. 9A) would then account for the observed decrease in cleft depth in narrower fins (Fig. 5D). On the other hand, the transition from a stretching tensile force at the tip to a contraction force as the fin model approaches its final length (A1, A2, A3, Fig. 9B) suggests a process through which mechanical cues from the fluid could induce the bifurcation of bony rays. Lower tensile stress acting on the narrower fin model (A3, B, Fig. 9B) offers a plausible explanation for the increased fraction of unbifurcated rays and enhanced distal shift observed in thinner regenerates (Fig. 6B). Our results on fins of different shapes (A2, C, D, Fig. 9) give interesting predictions on the forces acting during the regeneration of a typical zebrafish caudal fin. The simultaneous emergence of the bilobed shape together with the lengthening of the regenerate presumably leads to the emergence of stress distributions more complex than those measured on our stereotypical hydrofoils. Measurements on intermediate combinations of shape and length, both on actual fins as well as hydrofoils, would potentially bring more insights in the relation between forces, bifurcation and fin shape. In brief, our results suggest a consistent mechanical growth control scenario in which the dynamical map of dorso-ventral shear stress acting on the tip surface would regulate the growth rates profile across the rays and the bifurcation process.

Our study illustrates how the investigation of hydrodynamic stresses based on three-dimensional PTV experiments can provide important information about the mechanical interplay between a biological system and the surrounding fluid. We anticipate that hydrodynamic stresses will be integrated in a comprehensive growth model of the zebrafish caudal fin, as the most external and thus most upstream regulators. Indeed, our results suggest that the morphological fate of a bony ray in terms of growth rate and branching pattern is modulated by hydrodynamic forces. The works of (Murciano et al., 2002; Shibata et al., 2018) support this hypothesis as the grafted rays changed their bifurcation pattern or their length based on their new location, which could be explained by local differences in hydrodynamic stress. Understanding the role of the mechanical environment in the (re)establishment of positional information is of fundamental importance in the fields of regenerative medicine and tissue engineering.

To conclude, the experimental results presented in this work corroborate the hypothesis of a coupling between mechanical forces and morphogenesis, and support the idea that mechanical cues are transmitted from the surrounding fluid to the fin tissue to be integrated in the genetic cascade responsible for growth processes such as ray bifurcation and the orchestration of the overall fin shape. The simplicity of the regeneration experiments combined with stress measurements on hydrofoils, and the set of results presented here pave the way for future studies on the plasticity of fins in relation to their hydrodynamic environment.

## Acknowledgements

We thank V. Zimmermann for excellent technical assistance and for fish care; Sahil Puri, Siddhartha Verma, Flavio Noca and Petros Koumoutsakos for interdisciplinary discussions.

## Author contributions

Conceptualization: T.A-W, A.J., C.M.A, P.D. S.B.; Methodology: P.D, S.B. C.M.A., A.J.; Software, P.D and S.B; Investigation, P.D, S.B. D.P, C.P.; Formal Analysis: P.D., S.B., C.M.A.; Writing - original draft preparation: P.D., A.J. Writing - review and editing: S.B., T.A-W, C.M.A; Visualization: S.B.; Funding Acquisition, C.M.A and A.J.; Resources, C.M.A and A.J.; Project Management: T. AW.; Supervision, C.M.A and A.J.

---

**Suppl. Fig. S1.**
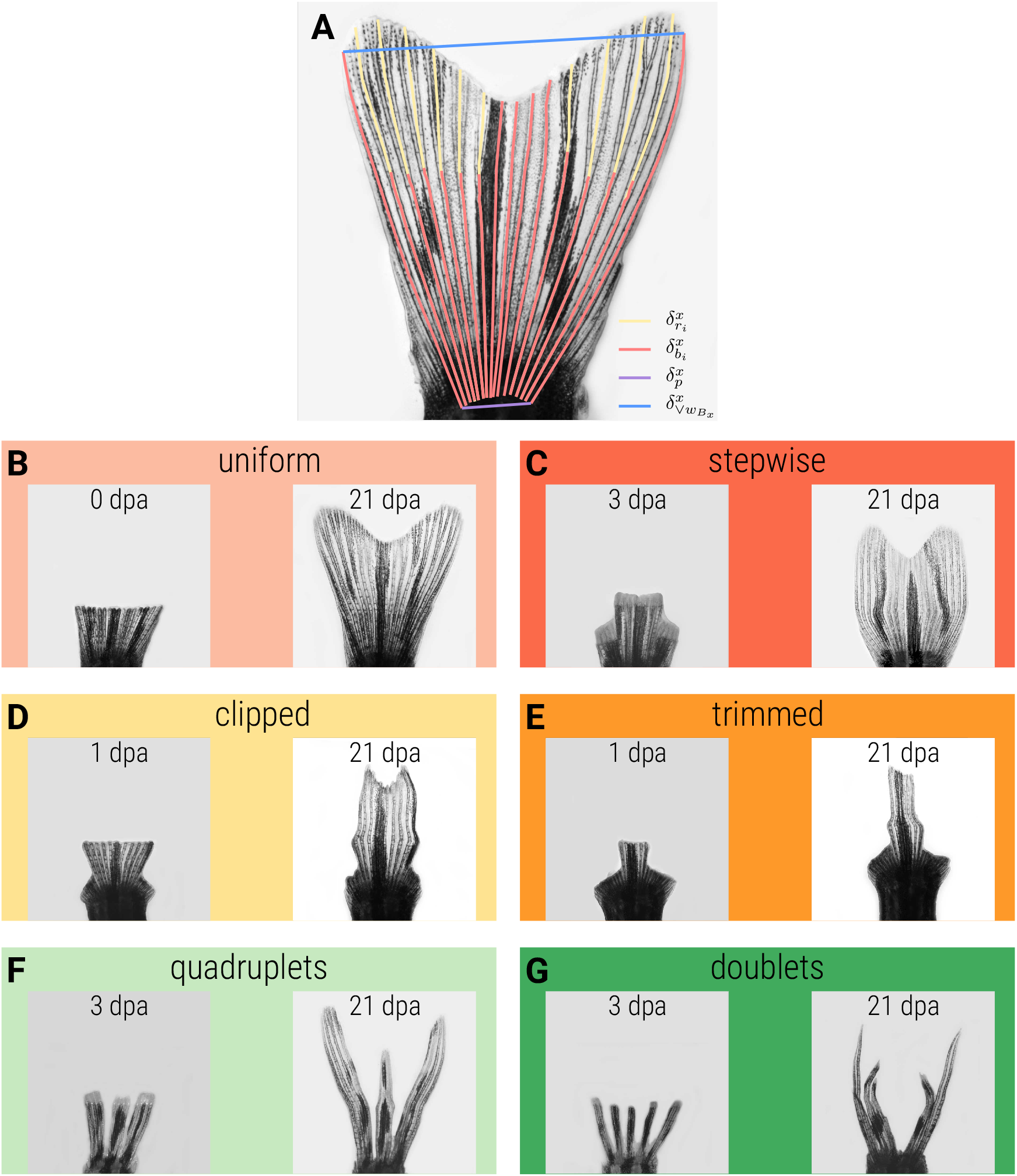
Representative images of fins with uneven amputations, as shown in schematic drawing in the main Fig. 3. **(A)** Morphological characterization of a fin. Ray length (δ_r_, yellow) is measured from the peduncle to the tip of the fin. Bifurcation length (δ_b_, red) is measured from the peduncle to the first bifurcation point. Peduncle width (δ_p_, purple) is measured between the two outermost rays. Fin width (δ_w_, blue) is measured between the two outermost nonablated rays, here corresponding to the two outermost rays. **(B-G)** Amputation strategies used in this study to challenge fin regeneration, overlaid with the corresponding naming convention. **(B, C)** Fins were allowed to naturally regenerate after surgery. **(D-G)** After the amputation procedure, selected rays were repeatedly ablated near the peduncle every 3 days for the duration of the experiment (21 days) to prevent their regeneration. In these situations the regenerated fins were lacking the ablated rays.

